# RPA phosphorylation regulates DNA resection

**DOI:** 10.1101/517771

**Authors:** Michael M. Soniat, Logan R. Myler, Tanya T. Paull, Ilya J. Finkelstein

## Abstract

Genetic recombination in all kingdoms of life initiates when helicases and nucleases process (resect) the free DNA ends to expose single-stranded (ss) DNA overhangs. Resection termination in bacteria is programmed by a DNA sequence but the mechanisms limiting resection in eukaryotes have remained elusive. Using single-molecule imaging of reconstituted human DNA repair factors, we identify a general mechanism that limits DNA resection. BLM helicase together with EXO1 and DNA2 nucleases catalyze kilobase-length DNA resection on nucleosome-coated DNA. The resulting ssDNA is rapidly bound by RPA, which is in turn phosphorylated as part of the DNA damage response (DDR). Remarkably, phosphorylated RPA (pRPA) inhibits DNA resection via regulation of BLM helicase. pRPA suppresses BLM initiation at DNA ends and promotes the intrinsic helicase strand-switching activity. These findings establish that pRPA is a critical regulator of DNA repair enzymes and provides a feedback loop between the DDR and DNA resection termination.

## Introduction

Double-strand DNA breaks (DSBs) are toxic DNA lesions that must be repaired rapidly and accurately(Vilenchik and Knudson, 2003, 2006). Incorrect repair of DSBs leads to oncogenic genome instability. Homologous recombination (HR) is a universally conserved DSB repair pathway that uses the information stored in an intact sister chromatid to repair the physical and genetic continuity of the broken genome(Jasin and Rothstein, 2013). In eukaryotes, homologous recombination is initiated by the MRE11-RAD50-NBS1 (MRN) complex, which rapidly localizes to DSBs in human cells (Lisby et al., 2004). MRN initiates HR by removing adducts from the DNA ends and loading the Bloom’s Syndrome helicase (BLM) along with Exonuclease 1 (EXO1)(Lisby et al., 2004; Myler and Finkelstein, 2016; Symington, 2016). An alternative pathway uses DNA2 helicase/nuclease instead of EXO1. BLM and EXO1 (or DNA2) cooperate generate segments of single-stranded DNA (ssDNA) by nucleolytically degrading (i.e., resect) one strand of the free DNA ends(Cejka et al., 2010; Daley et al., 2014; Mimitou and Symington, 2008; Nimonkar et al., 2008, 2011; Niu et al., 2010; Symington, 2016; Zhu et al., 2008).

All kingdoms of life use a combination of nucleases and helicases to catalyze DNA resection. In bacteria, the RecBCD/AddAB family of helicases and nucleases resect the free DNA ends(Dillingham and Kowalczykowski, 2008; Wigley, 2013). Resection is terminated via recognition of χ-sites—short GC-rich sequences (e.g., 5’-GCTGGTGG in *E. coli*) that are over-represented throughout the genome(Blattner et al., 1997; Smith, 2012; Spies et al., 2007). RecBCD/AddAB undergo a conformational rearrangement at χ that attenuates nuclease activity and loads RecA recombinase on the 3’-ssDNA overhang(Dillingham and Kowalczykowski, 2008). However, a χ-like sequence has not been identified in eukaryotes. The mechanisms that measure and terminate DNA resection are not fully understood. This is a critical question as eukaryotic DNA resection can proceed for tens of kilobases away from the DNA break when the free DNA ends lack homology to other genomic loci(Mimitou and Symington, 2008; Zhu et al., 2008).

Replication protein A (RPA) rapidly coats the ssDNA that is generated during DNA resection. RPA-ssDNA filaments activate the DNA damage response kinase ATR(Byun et al., 2005; Choi et al., 2010; Cortez et al., 2001; Zou and Elledge, 2003). ATR, together with ATM, CDK, and DNA-PKcs, phosphorylate RPA(Binz et al., 2004; Block et al., 2004; Liaw et al., 2011; Liu et al., 2012; Maréchal and Zou, 2015; Olson et al., 2006; Shiotani et al., 2013; Vassin et al., 2009). Though RPA phosphorylation is induced in response to DNA damage and is commonly used as readout a of DSB resection, cells expressing phosphomimetic RPA mutants have defects in DNA recombination and repair(Binz et al., 2003, 2004). Phosphorylation of RPA also modulates interactions between the N-terminus of RPA70 and other replication and repair proteins such as polymerase α, p53, MRN, and RAD51(Maréchal and Zou, 2015). However, the role of RPA phosphorylation during DNA resection and the functional significance of how RPA phosphorylation tunes protein-protein interactions in DSB repair remains unclear.

Here, we use single-molecule fluorescence imaging to establish that RPA phosphorylation is a critical regulator of eukaryotic resection on chromatin. BLM, in concert with RPA, stimulates processive resection by EXO1 and DNA2 nucleases. However, RPA32 phosphorylation inhibits DNA resection by altering the direct interaction between RPA70N and BLM. Phosphorylated RPA (pRPA) drastically slows both BLM/EXO1 and BLM/DNA2 resectosomes and stimulates BLM strand switching when the nuclease is omitted from the reaction. Moreover, BLM/EXO1 and BLM/DNA2 can resect past nucleosomes in the presence of RPA but are blocked when pRPA is added to the reaction. Thus, phosphorylated RPA is a critical negative regulator of DNA resection and other processes that involve BLM helicase.

## Results

### Mechanism of BLM/EXO1-mediated DNA resection

We established a single-molecule DNA curtains assay to image the role of RPA during DNA resection (**Figure 1**). In this assay, BLM helicase and the nucleases EXO1 or DNA2 are imaged on 48.5 kb-long DNA molecules that are organized on the surface of a lipid-coated microfluidic flowcell(Gallardo et al., 2015; Soniat et al., 2017). One end of the DNA substrate is biotinylated and tethered to streptavidin deposited on the lipid bilayer. The second DNA end terminates in a 3’-78 nucleotide (nt) overhang that mimics initial pre-processing of DNA breaks by the MRN/CtIP nuclease complex(Cannavo and Cejka, 2014; Myler et al., 2016, 2017; Paull, 1998; Shibata et al., 2014). In addition to tethering the DNA molecules, the fluid lipid bilayer also provides an inert surface for single-molecule experiments (**Figure 1A**). For fluorescent imaging, recombinant BLM was labeled with an anti-FLAG (when with EXO1) or anti-HA antibody (when with DNA2) conjugated to a fluorescent nanoparticle (quantum dot; QD) (**Figure S1A**). Biotinylated EXO1 was coupled to a streptavidin-conjugated QD that emits in a spectrally distinct fluorescent channel (**Figures 1B and S1A**)(Myler et al., 2016, 2017). DNA2 was imaged by an anti-FLAG antibody conjugated QD. This assay permits real-time imaging of both enzymes as they process the DNA substrate.

**Figure 1.**
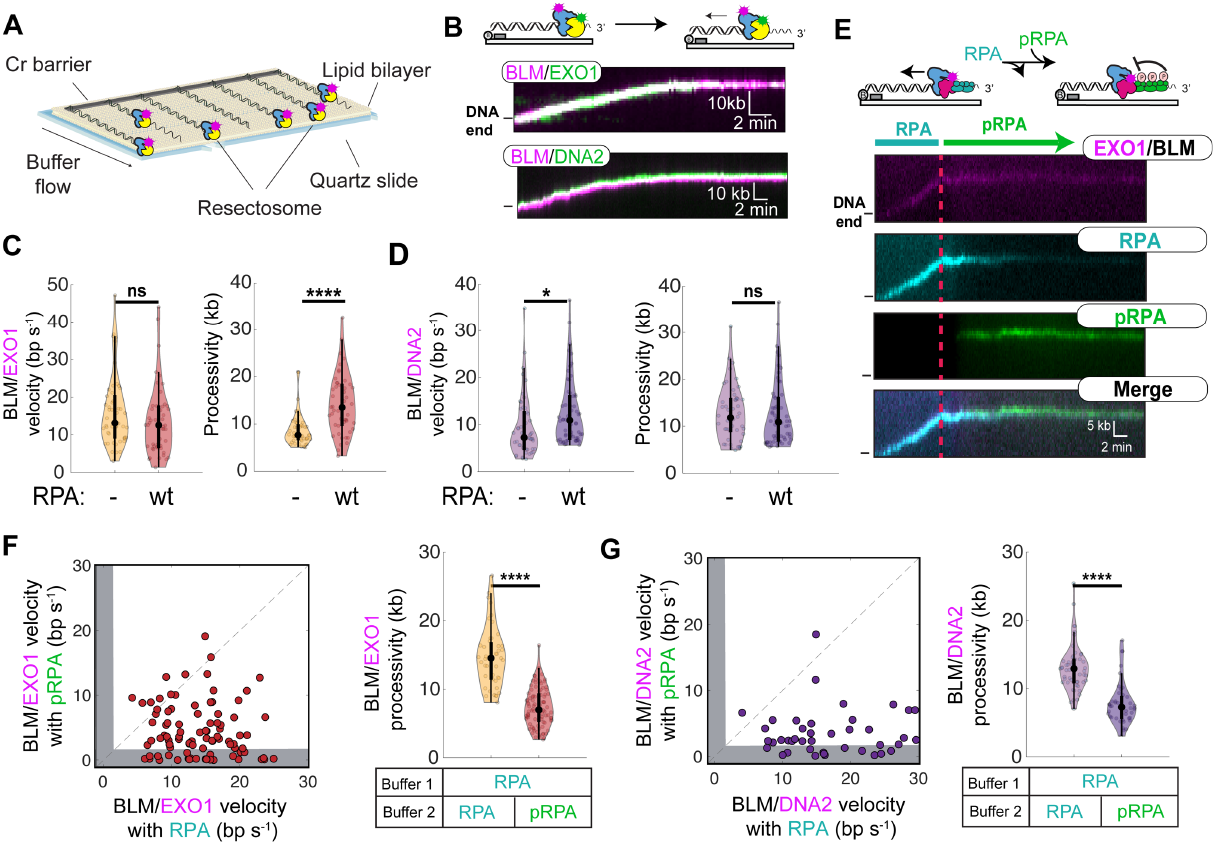
RPA regulates DNA resection. (**A**) Schematic of the single-molecule DNA resection assay. DNA substrates are organized at microfabricated barriers for single-molecule imaging. (**B**) Representative kymographs of BLM (magenta) and EXO1 (green) or BLM (magenta) and DNA2 (green) resecting DNA. (**C**) BLM/EXO1 and (**D**) BLM/DNA2 velocities and processivities with and without RPA. Dot in the violin plots represents the median and black bars indicate the interquartile range (thick black bar in the center) and 95% confidence intervals (thin black bars) of the distributions. (**E**) Kymograph of BLM and EXO1 (magenta) resecting DNA in the presence of 1 nM RPA-RFP (cyan) for 10 minutes before switching to pRPA-GFP (green) for 40 minutes. (**F-G**) Velocities (left) and processivities (right) of individual BLM/EXO1 (**F**) and BLM/DNA2 (**G**) complexes before and after switching from RPA to pRPA. Gray bars: molecules that stopped resection within our experimental uncertainty. Dashed line is shown as a reference with a slope of m=1. (p>0.05; ns, p<0.05; *,p<0.05; **, p<0.01; ***, p<0.001; ****, p<0.0001).

As EXO1 is the major DNA resection nuclease in human cells, we first assayed the BLM/EXO1 complex at the single-molecule level(Bolderson et al., 2010; Gravel et al., 2008; Myler et al., 2016; Nimonkar et al., 2011; Tomimatsu et al., 2012, 2012; Zhu et al., 2008). BLM and EXO1 physically interact in the absence of DNA and co-localize to free DNA ends in the single-molecule assay (**Figures 1B and S1B,C**)(Nimonkar et al., 2008, 2011). To monitor DNA resection, we pre-loaded the BLM/EXO1 complex on DNA ends and initiated translocation by supplementing the reaction buffer with 2 mM MgCl2 and 1 mM ATP. Under these conditions, ~91% (N=50/55) of the BLM/EXO1 complexes translocated at least ~1 kb away from the free DNA end within 129 ± 9 s of ATP being introduced, signaling initiation of DNA resection (**Figure 1B**). We had previously observed that EXO1 moves processively only after a long initiation time (750 ± 380 s; mean ± st. dev.)(Myler et al., 2016). These results indicate that BLM stimulates EXO1 initiation on DNA, possibly by opening the DNA duplex and aiding EXO1’s loading on the 5’-end, or via conformational regulation of a putative EXO1 autoinhibition domain(Orans et al., 2011; Shi et al., 2017; Warren et al., 2007).

BLM increased both the velocity and processivity of EXO1-catalyzed DNA resection. Compared to EXO1 alone, the velocity of the BLM/EXO1-complex increased 1.4-fold (16 ± 9 bp s^−1^; p<0.0001, N=50). The processivity was increased 2-fold (9 ± 3 kb; p<0.0001, N=50), consistent with BLM’s previously-reported role in stimulating DNA resection both *in vitro* and *in vivo* (**Figure 1C**)(Mimitou and Symington, 2008; Nimonkar et al., 2008, 2011; Niu et al., 2010; Yang et al., 2013). We also analyzed the velocity and processivity of EXO1 in complex with the helicase-dead BLM(K695A), which was reported to stimulate EXO1 via an unknown mechanism (**Figures S1D,E**)(Nimonkar et al., 2008). Helicase-dead BLM(K695A) did not change the velocity of the complex but increased the processivity to the same levels as wild type (wt) BLM (**Figures S1D,E**). Helicase-dead BLM can thus act as a processivity factor for EXO1 (also see below). BLM/EXO1 complexes terminated resection when both enzymes remaining co-localized but stopped moving (stalled) on DNA (**Figure 1B**). This observation suggests that one or both subunits can disengage their motors from processive DNA translocation in the stalled complex. In sum, BLM stimulates long-range DNA resection via both helicase-dependent and independent mechanisms that open the DNA substrate and retain EXO1 on DNA.

BLM also stimulated DNA2 resection at DNA ends (**Figure 1B**). Consistent with prior reports, DNA2 was inactive on 3’-ssDNA overhangs (**Figure S1F**)(Cejka et al., 2010; Nimonkar et al., 2011; Zhou et al., 2015). In contrast, 93% (N=50/54) of BLM/DNA2 complexes resected within 70 ± 4 seconds of ATP being introduced for 13 ± 6 kb at a rate of 9 ± 6 bp s^−1^ (**Figure 1D**). The addition of either helicase-dead BLM(K695A) or nuclease-dead DNA2(D277A) ablated DNA resection by the entire complex (**Figure S1F,G**). Similar to BLM/EXO1, BLM/DNA2 complexes terminated resection with both enzymes remaining co-localized in the stalled complex.

RPA rapidly coats all cellular ssDNA and also regulates BLM, EXO1, and DNA2(Brosh et al., 2000; Cannavo et al., 2013; Chen et al., 2013; Myler et al., 2016; Nicolette et al., 2010; Nimonkar et al., 2011; Sturzenegger et al., 2014; Yang et al., 2013; Yodh et al., 2009; Zhou et al., 2015). RPA physically interacts with BLM and stimulates its helicase activity(Brosh et al., 2000; Yodh et al., 2009). In contrast, RPA terminates processive EXO1 translocation by stripping the nuclease from DNA(Brosh et al., 2000; Genschel and Modrich, 2003; Genschel et al., 2002; Myler et al., 2016). RPA also directs DNA2’s nuclease activity to target the 5’ strand, leaving the 3’ end intact(Masuda-Sasa et al., 2006; Zhou et al., 2015). We therefore set out to define how RPA regulates DNA resection in the context of the BLM/EXO1 and BLM/DNA2 complexes.

The BLM/EXO1 and BLM/DNA2 resection reactions were supplemented with 1 nM RPA or RPA-GFP to visualize the ssDNA resection product. The intensity of the RPA-GFP signal increased proportionally with the distance traveled by BLM/EXO1 or BLM/DNA2, indicating that ssDNA is continuously generated during DNA resection (**Figure S1H**). With RPA, BLM/EXO1 complexes initiated long-range DNA resection within 56 ± 6 s (N=50) of ATP entering the flowcell. BLM/EXO1 velocity was statistically indistinguishable from that without RPA, but resection was 1.6-fold more processive (14 ± 6 kb; N=50) (**Figure 1C**). RPA also stimulated BLM/DNA2 via a slightly different mechanism. Nearly all (92%; N=60/65) BLM/DNA2 complexes initiated resection within 32 ± 3 s (N=60) of ATP entering the flowcell. However, RPA increased BLM/DNA2 velocity 1.4-fold (13 ± 7 bp s^−1^; N=60), but did not alter the resection processivity (13 ± 4 kb; N=60) (**Figure 1D**). The subtly different effects of RPA on the two complexes likely reflects the inhibitory effects of RPA on EXO1, but not on BLM or DNA2. Taken together, these data demonstrate that RPA stimulates initiation of the BLM/EXO1 and BLM/DNA2 resectosomes and promotes rapid, processive DNA resection.

### Phosphorylated RPA (pRPA) inhibits DNA resection

Phosphorylation of RPA induces conformational changes within the RPA70 subunit(Binz et al., 2003, 2004; Maréchal and Zou, 2015). We reasoned that these conformational rearrangements may alter how pRPA interacts with the BLM/EXO1 and BLM/DNA2 resectosomes to regulate DNA resection. To test this hypothesis, pRPA and fluorescent pRPA-GFP were prepared by incubating the recombinant RPA (overexpressed in *E. coli*) with SV40 replication-competent human cell extracts (**Figure S2A,B**)(Fotedar and Roberts, 1992; Stillman and Gluzman, 1985).

In the cell, pRPA is initially absent but begins to accumulate as a result of DNA resection(Maréchal and Zou, 2015). To recapitulate pRPA accumulation *in vitro*, DNA resection was imaged for 10 minutes with RPA-RFP in the flowcell and then switched to a buffer containing pRPA-GFP for another 40 minutes. This three-color single-molecule experiment allowed simultaneous observation of BLM/EXO1 (via an EXO1 fluorescent label), RPA-RFP, and pRPA-GFP (**Figure 1E**). As expected, pRPA rapidly replaced wt RPA on the ssDNA(Gibb et al., 2014). Strikingly, addition of pRPA caused 33% (N = 30/90) of BLM/EXO1 molecules to stop resecting the DNA (**Figure 1F**). The remaining 67% of the molecules moved at a ~2-fold lower velocity (6 ± 4 bp s^−1^; N=60/90) than with unphosphorylated RPA. BLM/EXO1 resection processivity was also reduced ~3-fold after addition of pRPA (**Figure 1F**). Similarly, upon addition of pRPA, 30% (N = 15/50) of BLM/DNA2 molecules stopped resecting the DNA with the remaining 70% of the molecules moving ~3-fold slower (6 ± 5 bp s^−1^; N=35/50) than with unphosphorylated RPA (**Figure 1G**). BLM/DNA2 resection processivity was also reduced ~3-fold after switching from RPA to pRPA (**Figure 1G**). Control experiments where wt unphosphorylated RPA was present in both resection buffers showed no change in velocity or processivity (**Figure 1F,G**).

We reasoned that RPA70N may be critical for regulating DNA resection because it physically interacts with both BLM and the phosphorylated N-terminus of RPA32(Brosh et al., 2000; Kang et al., 2018; Maréchal and Zou, 2015) (**Figure 2A**). At least two acidic patches within the N-terminus of BLM interact with RPA70N (**Figure 2B**)(Kang et al., 2018). To test the importance of the BLM-RPA70N interaction in regulating DNA resection, we assayed DNA resection in the presence of RPAΔN, which lacks the first 168 N-terminal residues of the RPA70 subunit. RPAΔN inhibited both BLM/EXO1 and BLM/DNA2 processivity and velocity (**Figure 2C-E**). This result is consistent with a second experiment where the RPA70N inhibitor 3,3’,5,5’-tetraiodothyroacetic acid was included in the resection buffer(Kang et al., 2018). This compound binds within the basic cleft of RNA70N and blocks interactions with ATRIP and BLM(Kang et al., 2018; Souza-Fagundes et al., 2012). In the absence of RPA, adding 100 μM of the inhibitor had no effect on BLM/EXO1 or BLM/DNA2 resection (**Figure S2D,E**). However, the compound inhibited both BLM/EXO1 and BLM/DNA2 velocity ~3-fold compared to resection with wt RPA. In addition, the compound resulted in a 3-fold decrease in processivity for both BLM/EXO1 and BLM/DNA2, respectively when used in concert with wt RPA. (**Figure 2D,E**). Lastly, we performed resection assays with *S. cerevisiae* RPA (yRPA) because the yeast RPA70N is only 20% identical with human RPA70N. Most of the RPA70N residues implicated in BLM interaction vary between the human and yeast RPA70 variants(Kang et al., 2018) (**Figure S2F**). Both the processivity and velocity of BLM/EXO1 and BLM/DNA2 were decreased in the presence of yRPA similar to pRPA.

**Figure 2.**
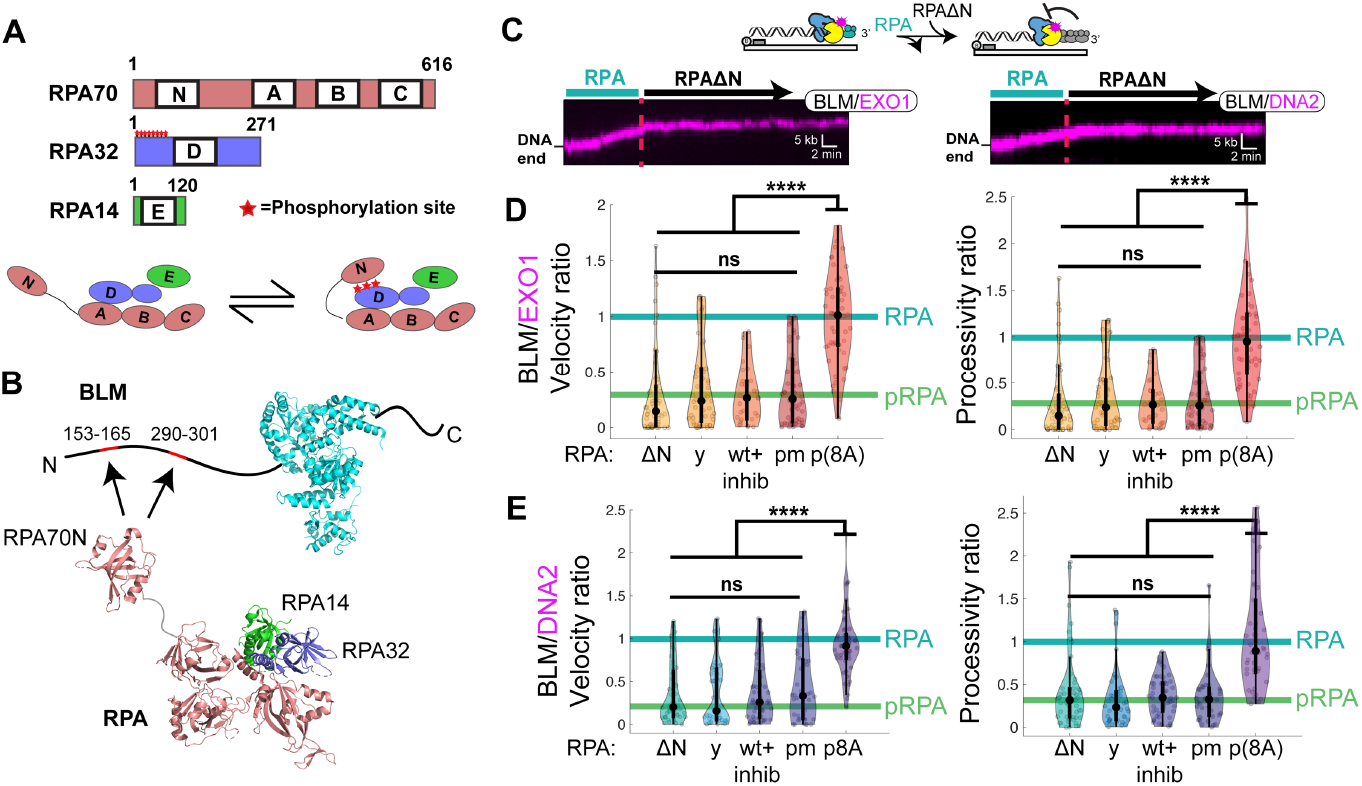
RPA70N-BLM interactions inhibit DNA resection. (**A**) RPA is phosphorylated on the RPA32 subunit at residues Ser4, Ser8, Ser11, Ser12, Ser13, Thr21, Ser23, Ser29, and Ser33. Bottom: RPA32 phosphorylation induces physical interactions with the N-terminus of RPA70 (RPA70N)(Binz et al., 2003, 2004; Maréchal and Zou, 2015). (**B**) BLM (PDB:4CGZ) interacts with RPA70N (PDB:4IPC) on RPA (PDB:4GOP) via at least two N-terminal acidic patches(Fan and Pavletich, 2012; Feldkamp et al., 2013; Kang et al., 2018; Newman et al., 2015). (**C**) Kymographs of BLM/EXO1 (left) and BLM/DNA2 (right) resecting DNA in the presence of 1 nM RPA for 10 minutes before switching to RPA AN for 40 minutes. (D) Ratios BLM/EXO1 and (**E**) BLM/DNA2 velocities (left) and processivities (right) before and after buffer switch with RPA variants: RPA AN (AN), yRPA (y), wt RPA plus RPA70 inhibitor (wt+inhib), phosphomimetic RPA (pmRPA), and phosphorylated RPA(8A) (p(8A)). Both velocity and processivity are compared to the ratios of wt RPA (cyan bar) and pRPA (green bar) from Figure 1F,G. (not significant; ns, p<0.05; *,p<0.05; **, p<0.01; ***, p<0.001; ****, p<0.0001).

pRPA prepared from 293T cell extracts is likely a heterogeneous ensemble of phosphorylation states. We therefore also purified phosphomimetic RPA (pmRPA) and phosphoblocking RPA(8A) (**Figure S2C**). pmRPA substitutes the RPA32 residues at positions 8, 11-13, 21, 23, 29, and 33 for asparagines and recapitulates RPA phosphorylation phenotypes *in vitro* and *in* vivo(Binz et al., 2003; Lee et al., 2010). RPA(8A) has these eight residues mutated to alanines, preventing their phosphorylation in 293T cell extracts. Switching from wt RPA to pmRPA drastically reduced the BLM/EXO1 and BLM/DNA2 velocity and processivity to levels that were statistically indistinguishable from resection with pRPA (**Figure 2D,E**). In contrast, switching from wt RPA to pRPA(8A)—RPA(8A) incubated in 293T cell extracts—did not inhibit DNA resection (**Figure 2D,E**). As RPA regulates all three enzymes, we also assayed the effects of pRPA on EXO1 and DNA2 in the absence of BLM. EXO1 alone was rapidly stripped from DNA by pRPA, consistent with our prior study showing how RPA inhibits EXO1 (**Figure S2G**)(Myler et al., 2016). Moreover, DNA2 alone did not initiate translocation from 3’-ssDNA ends with either RPA or pRPA (**Figure S2H**). Thus, we inferred that RPA interacts with BLM to regulate DNA resection (see below).

### RPA70N regulates BLM strand-switching

To understand how RPA regulates BLM, we first characterized the helicase on DNA curtains (**Figure 3A**). In the absence of RPA, 30% (N=90/300) of BLM molecules initiated long-range translocation 366 ± 26 s (N=90) after 1 mM ATP entered the flowcell (**Figure 3A**). As expected, helicase-dead BLM(K695A) remained stationary on DNA ends with 1 mM ATP in the imaging buffer (**Figure 3A and S3A**). Since BLM hasn’t been characterized on long DNA substrates at the single-molecule level, we next measured the ATP concentration-dependent velocity and processivity (**Figure 3B**). The results were well-described by a Michaelis–Menten fit with a K_m_ of 0.3 ± 0.2 mM ATP. The maximal velocity, V_max_, was 22 ± 4 bp s^−1^ with a maximal processivity of 15 ± 3 kb (**Figure 3B**). In all subsequent experiments, fluorescent BLM was monitored in the presence of 1 mM ATP with 1 nM of RPA-GFP (or wt RPA) in the imaging buffer (**Figure 3C and S3A**).

**Figure 3.**
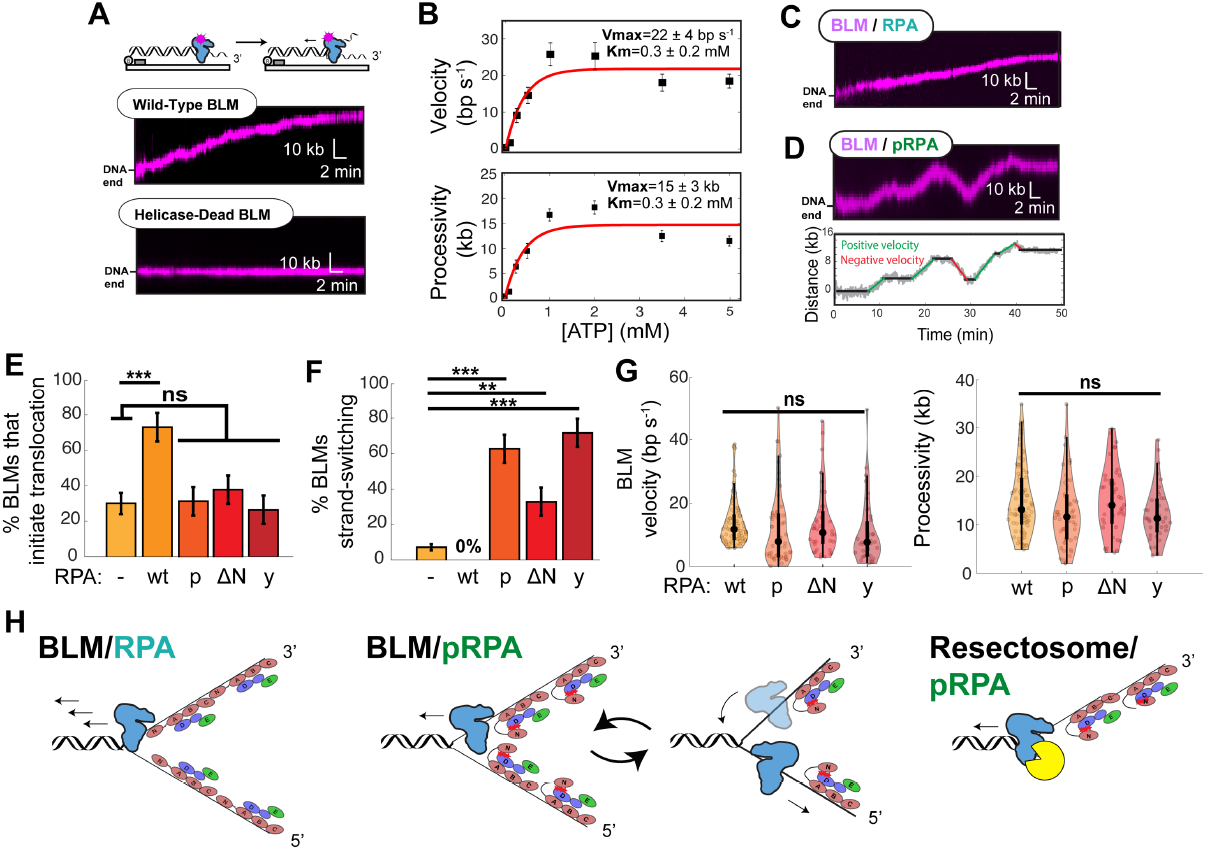
Phosphorylated RPA triggers BLM strand-switching. (**A**) BLM is a processive helicase whereas the helicase-dead BLM(K695A) does not move on DNA. (**B**) ATP concentration-dependent BLM velocity (top) and processivity (bottom) fit to Michaelis-Menten kinetics (red). Error bars: SEM. Fit parameters and 95% CI are indicated. (**C**) Kymograph of BLM (magenta) in the presence of 1 nM RPA or (**D**) with pRPA. Bottom: particle-tracking trace of the kymograph above highlighting pausing (black) and strand-switching (red, green) segments of the trajectory. (**E**) RPA stimulates BLM helicase initiation. This requires BLM-RPA70N interactions. RPA variants: wt RPA (wt), pRPA (p), RPA AN (AN), and yRPA (y). Error bars: S.D. as determined by bootstrap analysis. (**F**) BLM strand-switching is suppressed via BLM-RPA70N interactions. (**G**) Distribution of BLM velocities and processivities with RPA variants. (**H**) Model of RPA and pRPA regulation of BLM helicase and BLM in concert with either EXO1 or DNA2 nuclease. (ns, p>0.05; *,p<0.05; **, p<0.01; ***, p<0.001; ****, p<0.0001).

BLM helicase activity resulted in RPA-GFP accumulation on ssDNA which co-localized with the translocating BLM after ATP was added to the flowcell (**Figure S3B**). The velocity and processivity of fluorescent BLM with RPA-GFP were statistically indistinguishable from BLM without a fluorescent label (**Figure S3A**). wt RPA also increased the number of BLM molecules that initiated helicase activity (75%; N=80/107) and shortened the initiation time to 195 ± 14 s (N=80) (**Figure 3E and S3C**). With RPA, BLM’s velocity was reduced ~1.7-fold (15 ± 10 bp s^−1^; N=80), to the level seen with BLM/EXO1 (see **Figure 1C**). The processivity was statistically indistinguishable from experiments without any RPA (15 ± 7 kb; N=80) (**Figure 3G**). Thus, RPA stimulates BLM helicase initiation at ss/dsDNA junctions but reduces the helicase velocity.

BLM translocation was qualitatively different with pRPA, RPAΔN, and yRPA. These RPA variants failed to stimulate helicase initiation (**Figure 3E**). More strikingly, all three variants—but not wt RPA—induced the intrinsic strand-switching activity previously reported for BLM and other RecQ helicases (**Figure 3D,F and S3D**)(Chen and Brill, 2014; Harami et al., 2017; Klaue et al., 2013; Wang et al., 2015; Yodh et al., 2009). Strand-switching refers to BLM alternating between translocation on either the Watson or Crick ssDNA strands(Wang et al., 2015; Yodh et al., 2009). Prior smFRET observations of strand switching used short DNA oligos and those switching events were below the resolution of the DNA curtain assay(Wang et al., 2015; Yodh et al., 2009). Here, we observed several kb-long bursts of directional BLM translocation away from the ss/dsDNA junction (**Figure 3D**).

We quantified switches in translocation direction that were >2 kb, well above the ~500 bp resolution of the DNA curtains assay (**Figure 3F**). Without RPA, 8% (N=7/90) of BLM trajectories exhibited strand-switching. RPA completely suppressed such long-range strand-switching events (observed in 0% of 80 trajectories). In contrast, BLM strand switching was drastically increased with RPAΔN, yRPA, wt RPA plus the RPA70N inhibitor, and pRPA (**Figure 3F**). BLM switched strands once or twice per trajectory, followed by bursts of 2-4 kb-long processive segments (**Figure S3F**). To determine the effect of RPA variants on the overall processivity and velocity of BLM, we compared the total distance and average velocity of each directional change in BLM trajectories with the indicated RPA variants (**Figure 3G**). The results showed that although BLM strand-switches with pRPA, yRPA, and RPAΔN, the velocity and processivity are statistically indistinguishable from those with RPA after accounting for strand-switching. We thus conclude that RPA70N stimulates BLM helicase initiation and suppresses BLM’s intrinsic strand-switching activity (**Figure 3H**). Phosphorylation of RPA increases BLM strand switching, which may be especially important at reversing stalled replication forks or during branch migration and dissolution of Holliday junctions(Bizard and Hickson, 2014; Machwe et al., 2006; Ralf et al., 2006; Wu and Hickson, 2003). Moreover, BLM cannot switch strands in the context of the resection apparatus as one of the two ssDNA strands is degraded by either EXO1 or DNA2 nucleases (**Figure 3H, right**). During resection, BLM likely pauses translocation or stalls completely, leading to the reduced resection velocity and processivity that we observed with pRPA (**Figure 1**).

### Phosphorylated RPA inhibits resection past nucleosomes

We next sought to determine whether nucleosomes may provide an additional barrier to DNA resection. DNA substrates with an average of 4 ± 1 human nucleosomes per DNA molecule were prepared via stepwise salt dialysis (**Figure S4A**)(Brown et al., 2016). For fluorescent imaging, H2A with an HA epitope tag was visualized via a fluorescent anti-HA antibody(Myler et al., 2017).

Fluorescent EXO1 alone cannot resect past a nucleosome; 100% (N=40) of the molecules stalled at the first nucleosome that they encountered (**Figure 4A,D**). These results are in agreement with a report that yeast Exo1 activity is abrogated on a nucleosomal substrate(Adkins et al., 2013). Fluorescent DNA2 alone did not encounter any nucleosomes due to its inability to initiate resection on 3’-ssDNA overhangs with RPA (**Figure S1F**)(Cejka et al., 2010; Nimonkar et al., 2011; Zhou et al., 2015). In contrast, BLM in the presence of RPA pushed nucleosomes for an average of 6 ± 4 kb with a velocity of 9 ± 5 bp s^−1^ (N=50) (**Figure 4B,D and S4B**). Nucleosome collisions reduced both the BLM helicase processivity and velocity ~2-fold in the presence of RPA relative to naked DNA. All trajectories terminated with BLM stalling after pushing a nucleosome without any apparent loss or disassembly of the histone octamer, as reported by the fluorescent H2A signal.

**Figure 4.**
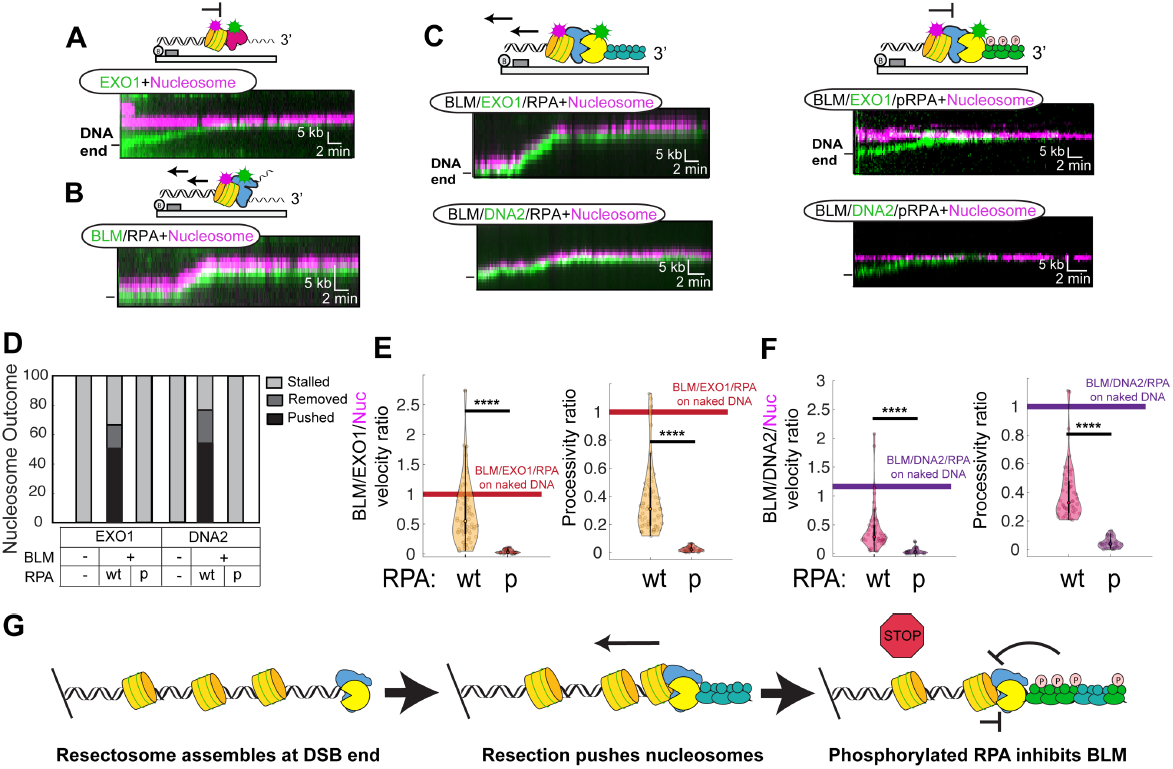
Nucleosomes and phosphorylated RPA terminate resection. (**A**) Representative kymograph showing EXO1 (green) stalling at a nucleosome (magenta). Nucleosome-coated DNA was reconstituted via stepwise salt dialysis of histone octamers onto the DNA substrate. (**B**) Representative kymograph showing that BLM pushes nucleosomes in the presence of wt RPA. (**C**) Representative kymograph showing that in the presence of wt RPA BLM/EXO1 and BLM/DNA2 (left) are able to move nucleosomes. However, in the presence of pRPA (right), both BLM/EXO1 and BLM/DNA2 stall at a nucleosome. (**D**) Outcomes of collisions for the indicated resectosome components. (**E**) Relative velocities and processivities of nucleosomes that encounter the indicated resectosome components. These values are normalized to the velocity and processivity measured for the same enzymes on naked DNA. In all cases, nucleosomes decrease both the velocity and processivity or completely stall resection. (ns, p>0.05; *,p<0.05; **, p<0.01; ***, p<0.001; ****, p<0.0001). (**F**) Model summarizing how phosphorylation of RPA acts as a negative regulator of DNA resection on chromatin.

Both BLM/EXO1 and BLM/DNA2 were able to resect past a nucleosome in the presence of wt RPA (**Figure 4C-F**). In >60% of such collisions, the nucleosome was pushed by the BLM/EXO1 (or DNA2) resectosome. Of these pushed nucleosomes, H2A signals were lost in 24% of BLM/EXO1 trajectories (N=16/68) and 30% of BLM/DNA2 trajectories (N=15/50). H2A loss can be due to disassembly of the histone octamer with a DNA-bound tetrasome or removal of the entire octamer from DNA. Nucleosome collisions reduced both the DNA resection processivity and velocity relative to naked DNA, showing that both resectosomes have difficulty pushing nucleosomes. Compared to BLM/EXO1 with RPA on naked DNA, the resection processivity was decreased ~3-fold (5 ± 4 kb; N=68) and the velocity was also decreased 2-fold (10 ± 8 bp s^−1^; N=68). Similarly, both the processivity and velocity of BLM/DNA2 were decreased 3-fold (5 ± 3 kb; 5 ± 4 bp s^−1^; N=50) compared to BLM/DNA2 in the presence of RPA on naked DNA (**Figure 4D-F**). Resection in the presence of pRPA showed drastically different nucleosome collision outcomes (**Figure 4C-F**). Both BLM/EXO1 and BLM/DNA2 were completely inhibited by the first nucleosome that the complexes encountered (N=40 and 35, respectively). Taken together, these results show that BLM assists both EXO1 and DNA2 to resect past nucleosome barriers. Nucleosomes remain associated with the DNA, as has been observed previously for the RecBCD complex(Eggleston et al., 1995; Finkelstein et al., 2010). Most surprisingly, RPA phosphorylation stalls the enzymes at the first nucleosome that they encounter on DNA.

## Discussion

The resection of free DNA ends initiates homologous recombination at double-stranded DNA breaks in all organisms, from bacteriophage to humans. Timely termination of DNA resection is critical for cell survival, as persistent over-resection of unrepaired DNA ends can lead to cell death (Chen et al., 2013; Rossiello et al., 2014; Zhu et al., 2008). In bacteria, resection termination is programmed at the DNA sequence level. For example, the *E. coli* RecBCD and *B. subtilis* AddAB helicase/nuclease complexes catalyze rapid and long-distance DNA resection until they encounter a species-specific GC-rich χ sequence(Dillingham and Kowalczykowski, 2008; Krajewski et al., 2014; Singleton et al., 2004; Wilkinson and Wigley, 2014). This sequence signals a conformational rearrangement that ultimately terminates DNA resection. The mechanism(s) regulating resection termination in eukaryotes do not appear to be encoded in a specific DNA sequence. One mechanism involves recruitment of the DNA helicase HELB by RPA, possibly to remove EXO1 from the DNA(Tkáč et al., 2016). In addition, EXO1 is degraded in a checkpoint-dependent manner via phosphorylation by ATR and ATM kinases(Bolderson et al., 2010; Cotta-Ramusino et al., 2005; El-Shemerly et al., 2008; Morin et al., 2008; Tomimatsu et al., 2017). Furthermore, DNA resection has recently been shown to be suppressed by the 53BP1 effector complex, shieldin(Dev et al., 2018; Ghezraoui et al., 2018; Gupta et al., 2018; Mirman et al., 2018; Noordermeer et al., 2018). However, a general mechanism for how DNA resection is terminated after initiation has not been identified.

Here, we show that a negative feedback between pRPA and BLM limits the extent of DNA resection on both naked and chromatinized DNA (**Figure 4G**). ssDNA that is generated during DNA resection via BLM/EXO1 or BLM/DNA2 resectosomes is rapidly coated by RPA. RPA stimulates long-range DNA resection, allowing helicase/nucleases to push individual nucleosomes for several kilobases on DNA. These nucleosomes are most likely removed by dedicated chromatin remodelers (e.g., SMARCAD1) if longer-range resection is required(Chen et al., 2012; Costelloe et al., 2012). ssDNA-bound RPA also recruits the ATR-ATRIP complex, which promotes phosphorylation of RPA(Liu et al., 2012; Olson et al., 2006; Shiotani et al., 2013; Vassin et al., 2009; Zou and Elledge, 2003). RPA32 is further phosphorylated on S4/S8 by DNA-PKcs which is a key NHEJ factor(Anantha et al., 2007; Liu et al., 2012; Maréchal and Zou, 2015). Furthermore, ATM has been shown to phosphorylate RPA32 at residues S4, S8, and S12 in response to DNA damage(Liu et al., 2012; Maréchal and Zou, 2015). The resulting accumulation of pRPA inhibits the processive DNA resection machinery on chromatin. Stalled resection machinery may then be removed by the DNA resection-associated helicase HELB.

We demonstrate that direct interactions between BLM and pRPA limit DNA resection. BLM is a processive helicase and RPA promotes BLM initiation at ss/ds junctions through an interaction with the N-terminus of the RPA70 subunit (**Figure 3H**). Phosphorylation of RPA abrogates or changes this interaction and decreases BLM’s ability to initiate at ss/ds junctions, in addition to promoting the intrinsic strand-switching activity of BLM. Regulation of BLM strand switching may be important during the later stages of HR (i.e., joint molecule dissolution), Holliday junction migration, resolution of ultra-fine bridges, and at stalled replication forks(Bachrati and Hickson, 2008; Bizard and Hickson, 2014; Croteau et al., 2014; Liu et al., 2014; Machwe et al., 2006; Ralf et al., 2006; Sarlós et al., 2018; Wu and Hickson, 2003). In the context of DNA resection, one of the two ssDNA strands are degraded by a nuclease, forcing BLM to slow down or stall.

This work underscores that BLM helicase activity and overall DNA resection is regulated by phosphorylation of the RPA heterotrimer. RPA70 is a critical interaction hub for multiple proteins involved in DNA repair (e.g., DNA2, MRN, FANCJ, RAD51), replication fork reversal (e.g., RAD52, WRN, SMARCAL1) and replication (e.g., RFC, Pol a primase, FACT)(Maréchal and Zou, 2015). RPA70 is also sequestered by phosphorylation at the N-terminus of RPA32(Binz et al., 2003, 2004; Maréchal and Zou, 2015). Our results indicate that phosphorylation-dependent changes in inter-subunit RPA interactions may be general regulators of DNA maintenance factors important for preserving genomic integrity.

## Materials and Methods

### Protein Cloning and Purification

Oligonucleotides were purchased from IDT. Human RPA, yeast RPA, pmRPA, RPA-GFP, and RPA truncation variants were purified from *E. coli* using a pET expression vector (Binz et al., 2003; Modesti, 2011; Myler et al., 2016). The phosphomimetic RPA contains the following amino acid substitutions in the RPA2 gene product: S8, S11, S12, S13, T21, S23, S29, S33 of RPA32 (Binz et al., 2003). Epitope-tagged human Exonuclease 1 (EXO1) was purified from insect cells (Myler et al., 2016).

A pFastBAC plasmid encoding the Bloom’s helicase gene (pIF458) was expressed in Sf21 insect cells infected using the Bac-to-Bac expression system (Life Tech.) (Yang et al., 2013). Cells were harvested 72 hours after infection, pelleted, frozen, and stored at −80°C. Cells were homogenized in buffer A containing 50 mM KH2PO4, pH 7.0, 500 mM KCl, 10% glycerol, 20 mM β-mercaptoethanol, 2.5 mM imidazole, and 250 mM phenylmethane sulfonyl fluoride (PMSF) in a Dounce homogenizer (Kimble Chase; Kontes) followed by sonication on ice. Insoluble material was pelleted for 1 hr at 35,000 rpm and supernatant was added to Ni-NTA resin (QIAGEN, 30410) in batch and rotated at 4°C for 1 hr. Ni-NTA resin was then spun at 3,000 rpm for 10 min and, washed 3x with buffer A. BLM was eluted with 15 mL of buffer B containing 50 mM KH_2_PO_4_, pH 7.0, 500 mM KCl, 10% glycerol, 20 mM β-mercaptoethanol, 250 mM Imidazole, and 250 mM PMSF. BLM was then incubated with Anti-FLAG M2 Affinity Gel (Sigma-Aldrich, A2220) at 4°C for 1 hr, washed with 3x with buffer C (25 mM Tris pH 8.0, 100 mM NaCl, 10% Glycerol, 1 mM DTT), and eluted with 5 mL of buffer C containing 0.1 mg/mL FLAG peptide (Sigma-Aldrich, F3290). BLM was further purified using a 1 mL HiTrap SP (GE Healthcare, 17115101) with a gradient from buffer C to buffer D (25 mM Tris pH 8.0, 1 M NaCl, 10% Glycerol, 1 mM DTT) and dialyzed overnight at 4°C in Buffer C.

For single-molecule fluorescent imaging, a 3xHA epitope tag was added onto the N-terminus of BLM via Q5 PCR mutagenesis. The 3xHA-BLM variant was purified using a similar protocol as FLAG-BLM, with the following minor modifications. After lysis and clarification in buffer A, the supernatant was purified using a 5 mL HisTrap HP column (GE Healthcare, 17524802) and eluted with buffer B. BLM was further purified using HiTrap SP (GE Healthcare, 17115101) and dialyzed overnight at 4°C in Buffer C.

A DNA2 pFastBAC plasmid encoding FLAG-DNA2 (pIF494) was generously provided by Jim Daley and expressed in Sf21 insect cells infected using the Bac-to-Bac expression system (Life Tech.) Cells were harvested 72 hours after infection, pelleted, frozen, and stored at −80°C. Cells were homogenized in 25 mM Tris·HCl pH 8.0, 100 mM NaCl, 10% (vol/vol) glycerol, 400 μL of PMSF (17 mg/mL), 20 mM β-mercaptoethanol in a Dounce homogenizer (Kimble Chase; Kontes) followed by sonication on ice. Insoluble material was pelleted for 1 hr at 35,000 rpm and supernatant was added to an anti-FLAG M2 Affinity Gel (Sigma-Aldrich, A2220) at 4°C for 1 hr, washed with 3x with buffer C (25 mM Tris pH 8.0, 100 mM NaCl, 10% Glycerol, 1 mM DTT), and eluted with 5 mL of buffer C containing 0.1 mg/mL FLAG peptide (Sigma-Aldrich, F3290). DNA2 was further purified using a 1 mL HiTrap SP (GE Healthcare, 17115101) with a gradient from buffer C to buffer D (25 mM Tris pH 8.0, 1 M NaCl, 10% Glycerol, 1 mM DTT) and dialyzed overnight at 4°C in Buffer C.

### Single Molecule Fluorescence Microscopy

#### Data acquisition

All single-molecule data were collected on a Nikon Ti-E microscope in a prism-TIRF configuration equipped with a Prior H117 motorized stage. Flowcells were loaded into a custom-designed stage insert incorporating a chip mount, fluidic interface, and heating element (Soniat et al., 2017). All experiments were maintained at 37°C by a combination of an objective heater (Bioptechs) and a custom-built stage-mounted heating block. The flowcell was illuminated with a 488 nm laser (Coherent) through a quartz prism (Tower Optical Co.). Data were collected with a 200 ms exposure, 1-second shutter (Vincent Associates) resulting in 3,600 frames in 1 hour, through a 60X water-immersion objective (1.2NA, Nikon), a 500 nm long-pass (Chroma) and a 638 nm dichroic beam splitter (Chroma), which allowed two-channel detection through two EMCCD cameras (Andor iXon DU897, cooled to −80°C). For three-color experiments, data was collected with 200 ms exposure, 2-second shutter, and a 561 nm dichroic beam splitter (Chroma), which allowed for detection of Quantum Dot-705 labeled EXO1, RPA-RFP, and pRPA-GFP. Images were collected using NIS-Elements software and saved in an uncompressed TIFF file format for later analysis (see below).

#### Data analysis

Fluorescent particles were tracked in ImageJ using a custom-written particle tracking script (available upon request). The resulting trajectories were analyzed in Matlab (Mathworks). Trajectories were used to calculate velocity and processivity for BLM or the BLM/EXO1 complex. EXO1 binding lifetimes were fit to a single exponential decay using a custom MATLAB script. Histograms of binding preferences for BLM and EXO1 on DNA were acquired by combining data from at least three flowcells for each experiment and fitting to a Gaussian curve using a custom script written in MATLAB.

To obtain Michaelis-Menten parameters for BLM translocation (V_max_ and K_m_), at least 35 individual BLM molecules were tracked at each ATP concentration at 37°C. The histogram of the velocity and processivity distributions were fit to Gaussian functions. The center of the fit is the reported value and the error-bars correspond to the standard deviation of the fits. After obtaining the mean velocity and processivity as a function of ATP, the data was then fit to a Michaelis–Menten curve.

DNA substrates for single-molecule studies contained a 78 nucleotide 3’ overhang. These were prepared by annealing oligonucleotides IF007 and LM003 to lambda phage DNA (Myler et al., 2016). For nucleosome reconstitution, the DNA was first ligated to the oligonucleotide handles and then concentrated using isopropanol precipitation and resuspension in TE buffer with high salt (10mM Tris-HCl pH 8.0, 1mM EDTA, 2M NaCl). Human octamers were reconstituted into nucleosomes at a nominal ratio of 1:100 (DNA:octamer) and dialyzed via a stepwise salt-dialysis using 1.5 M, 1.0 M, 0.8 M, 0.6 M, 0.4 M, and 0.2 M for 2 hrs each. The nucleosomes were then visualized on DNA by injecting a fluorescent antibody directed against a HA epitope tag on the H2A subunit.

#### Fluorescent protein labeling

FLAG-BLM (40 nM) was conjugated to Quantum Dots (QDs) by first pre-incubating a biotinylated anti-FLAG antibody (Sigma-Aldrich, F9291) with streptavidin QDs (Life Tech., Q10163MP for 705 and Q10103MP for 605) on ice for 10 minutes in 20 μL. Next, BLM was incubated with the anti-FLAG QDs at a ratio of 1:2 for an additional 10 minutes on ice, diluted with BSA buffer containing free biotin to 200 μL, and injected into the flowcell. 3xHA-BLM was labeled with anti-HA antibody (ICL Lab, RHGT-45A-Z) conjugated QDs on ice for 10 minutes prior to injection. EXO1 was conjugated to streptavidin QDs at a ratio of 1:2. Saturating biotin was added to the EXO1-QD conjugates to bind free streptavidin sites and the mixture was diluted to 200 μL prior to injecting EXO1 into the flowcell.

#### In vitro phosphorylation of human RPA

Purified human RPA was phosphorylated *in vitro* as described previously with a few minor modifications (Fotedar and Roberts, 1992; Stillman and Gluzman, 1985). Briefly, HEK293T cells were lysed in buffer A (25mM Tris-HCl pH 8, 100mM NaCl, 10% glycerol) by sonication. Next, the lysate was clarified by centrifugation at 20,000xg (Eppendorf Centrifuge 5424) for 10 minutes. The concentration of extract was measured via Bradford assay and adjusted to 10 mg mL^−1^. Next, the following components were combined in a 50μL solution with buffer B (40mM HEPES pH 7.5, 8mM MgCl2, 0.5mM DTT, 3mM ATP): 100 ng of pcDNA3 plasmid DNA (an SV40 replication origin containing plasmid, 10 mg mL^−1^), HEK293T extract (2-5 mg mL^−1^ final concentration), and 2μM purified RPA (final concentration: 400 nM). This reaction was incubated at 37°C for 2 hours. Phosphorylated RPA was purified from the extract using a 1 mL HiTrap Q. RPA phosphorylation was assayed by western blot using antibodies for pRPA at S4/S8 and S33 (Bethyl Laboratories).

### Pull-Down Assays

FLAG-BLM was incubated with biotinylated EXO1, two units of DNase I (NEB), and 20 ng of bovine serum albumin (BSA, Fisher Scientific) in Buffer A (25 mM Tris-HCl [pH 8.0], 100 mM NaCl, and 10% glycerol) for 30 minutes on ice. The samples were then added to a mixture of 100 ng BSA and 5 μL of streptavidin-coated paramagnetic beads (Dynabeads M-280, Life Tech.) for an additional 15 min incubation on ice. After three washes with 2 mg mL^−1^ BSA in Buffer A, proteins bound to the beads were resolved by 8% SDS-PAGE, followed by western blotting with anti-FLAG primary antibodies (Sigma-Aldrich, F1804), anti-mouse secondary antibodies (Rockland, RL-610-132-121), and streptavidin ATTO647N (Atto-Tec, AD 647N-65).

## End Matter

### Author Contributions and Notes

M.M.S. and L.R.M. prepared proteins and DNA samples. M.M.S. and L.R.M. conducted all single-molecule experiments and pull-down assays. T.T.P. provided critical reagents. T.T.P. and I.J.F. directed the project. M.M.S, L.R.M., and I.J.F. co-wrote the paper with input from all coauthors.

## Acknowledgments

We are indebted to our colleagues Jim Daley and Marc Wold for valuable reagents. We are grateful to members of the Finkelstein and Paull laboratories for useful discussions and for critically reading the manuscript. This work was supported by the National Institutes of Health (GM120554 to I.J.F.), the National Cancer Institute (CA092584 to I.J.F., CA212452 to L.R.M.), CPRIT (R1214 to I.J.F., RP110465 to T.T.P.), and the Welch Foundation (F-l808 to I.J.F.). I.J.F. is a CPRIT Scholar in Cancer Research. T.T.P. is an investigator of the Howard Hughes Medical Institute. Michael Soniat is supported by a Postdoctoral Fellowship, PF-17-169-01-DMC, from the American Cancer Society. The content is solely the responsibility of the authors and does not necessarily represent the official views of the National Institutes of Health.

