## Supplemental Information for "RPA phosphorylation regulates DNA resection"

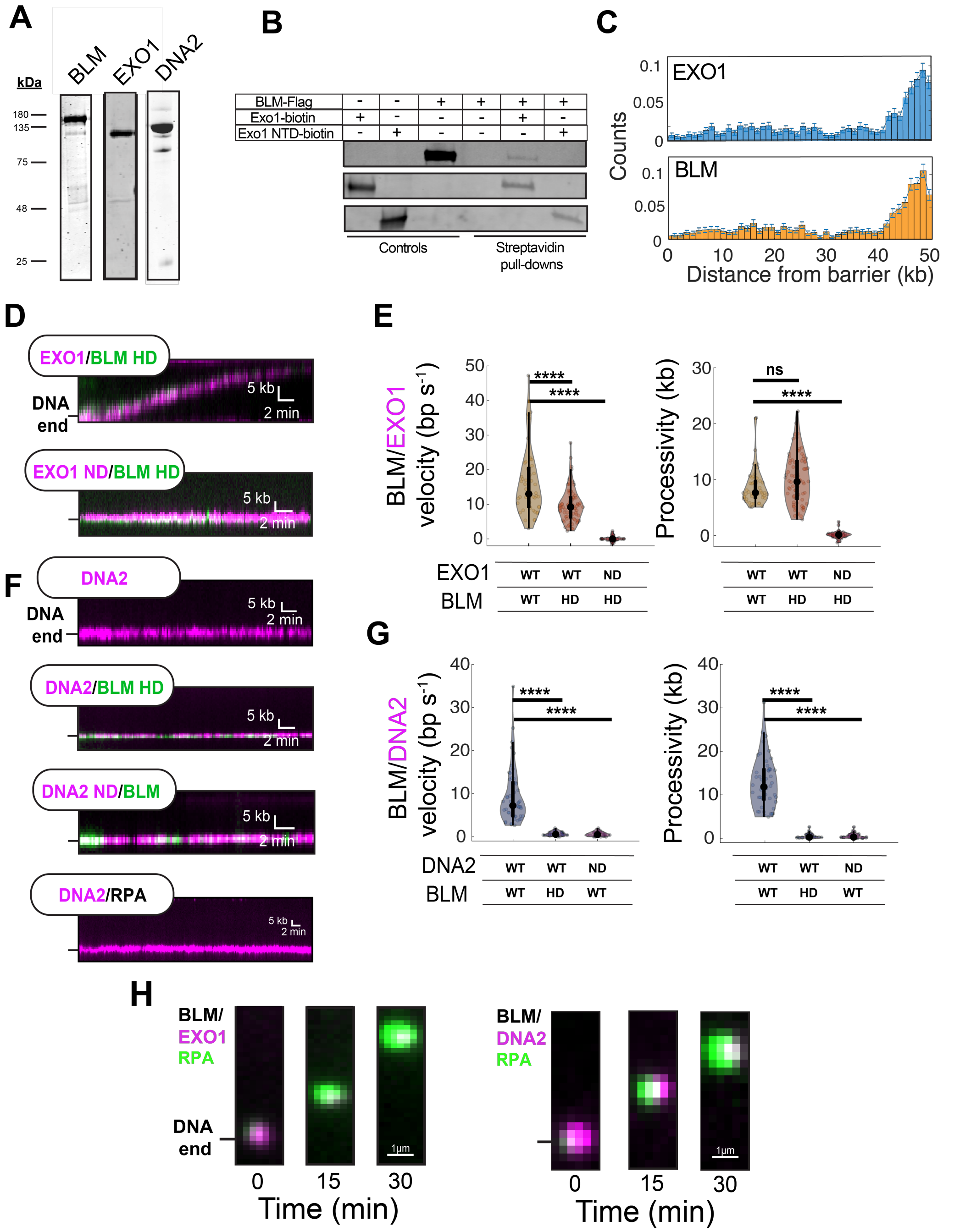


**Figure S1. Characterization of BLM/EXO1 and BLM/DNA2 resectosomes**

**(A)** SDS-PAGE gel of recombinantly expressed full-length BLM, EXO1, and DNA2. **(B)** Streptavidin pull-down assays showing BLM interacts with full-length EXO1 (lane 5) but not with the N-terminal catalytic domain (NTD; aa 1-352) of EXO1 (lane 6). **(C)** Histogram of BLM and

EXO1 binding along the DNA substrate. As expected, BLM and EXO1 co-localize at the free DNA ends. Error bars are the SD as determined by bootstrapping. **(D)** Representative kymograph showing that helicase-dead (HD) BLM(K695A)/EXO1 is still able to resect DNA. As expected, helicase-dead BLM in complex with nuclease-dead (ND) EXO1 (D78A/D173A) binds but does not leave the DNA ends. **(E)** Velocity and processivity of the indicated catalytically inactive BLM and EXO1 complexes. **(F)** Representative kymographs showing that helicase-dead (HD) BLM/DNA2 or BLM/DNA2(ND) is not able to resect DNA. In addition, neither DNA2 nor DNA2/RPA is able to resect DNA in the absence of BLM. **(G)** Velocity and processivity plots of helicase-dead (HD) BLM/DNA2 and BLM/DNA2 ND. **(H)** RPA-GFP intensity increased with BLM/EXO1 and BLM/DNA2 resection due to accumulation of ssDNA.

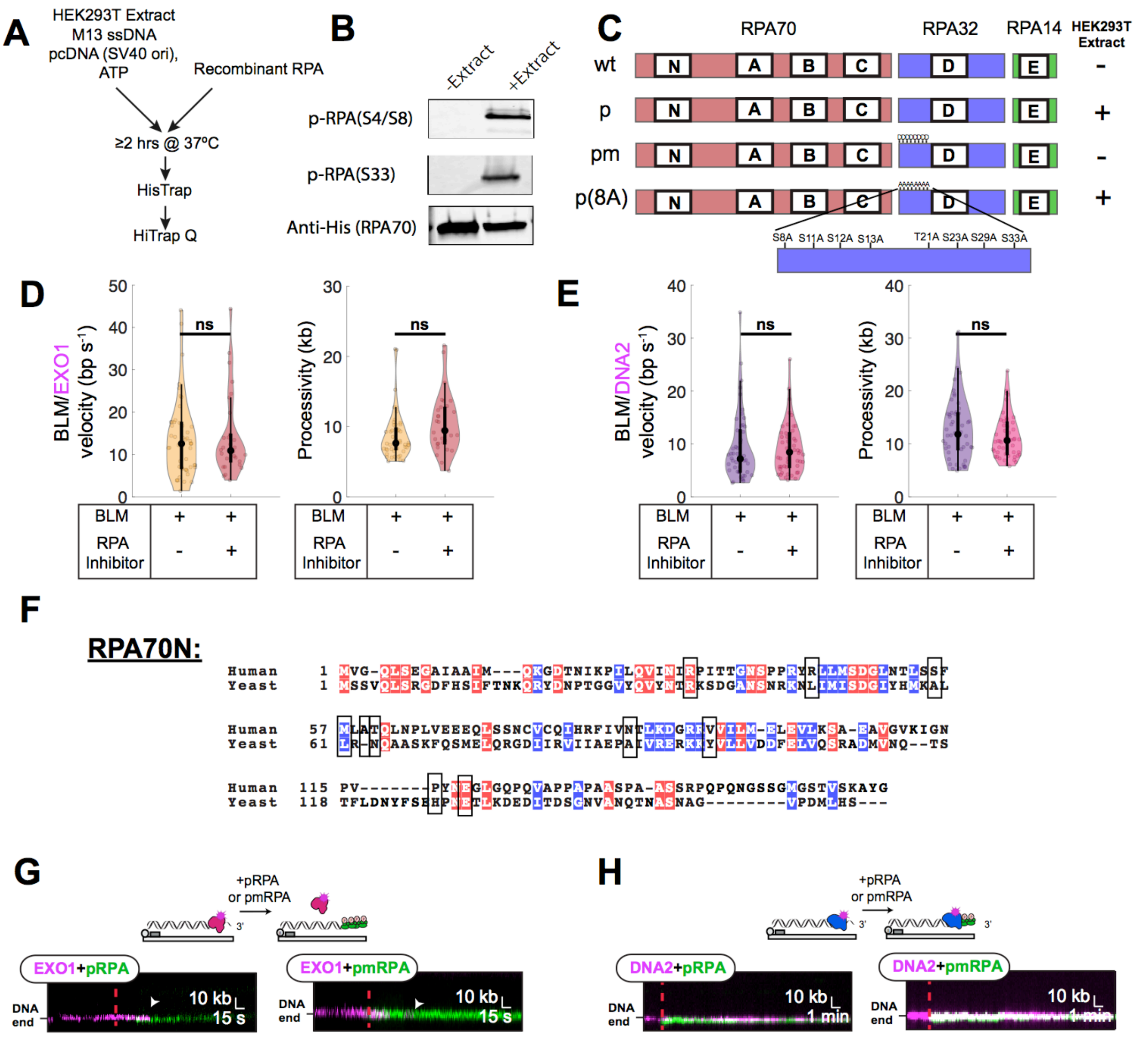


**Figure S2. Characterization of RPA70N interaction with resectosomes**

**(A)** Phosphorylation and purification of pRPA **(B)** Western blot monitoring pS4/S8 and pS33 of purified phosphorylated RPA with and without the addition of HEK293T extract. **(C)** Schematic of RPA variants used in current study. **(D-E)** Velocity and processivity distributions of BLM/EXO1 **(D)** and BLM/DNA2 **(E)** with and without RPA70N inhibitor 3,3',5,5'-tetraiodothyroacetic acid showing RPA inhibitor does not inhibit resection in absence of RPA. **(F)** Sequence alignment generated by CLUSTALW2 of RPA70N from human and yeast

(*Saccharomyces cerevisiae*)(Larkin et al., 2007). Homologous regions are box-shaded red (identical amino acid residues) and blue (conserved amino acid substitutions). Boxes represent residues important for BLM-RPA70N interactions(Kang et al., 2018). **(G)** Representative kymographs showing EXO1 is rapidly stripped from DNA in the presence of pRPA and pmRPA, as we had observed for wt RPA(Myler et al., 2016). White arrows indicate EXO1 dissociation. **(H)** Representative kymographs showing DNA2 remains on DNA but does not move with pRPA and pmRPA.

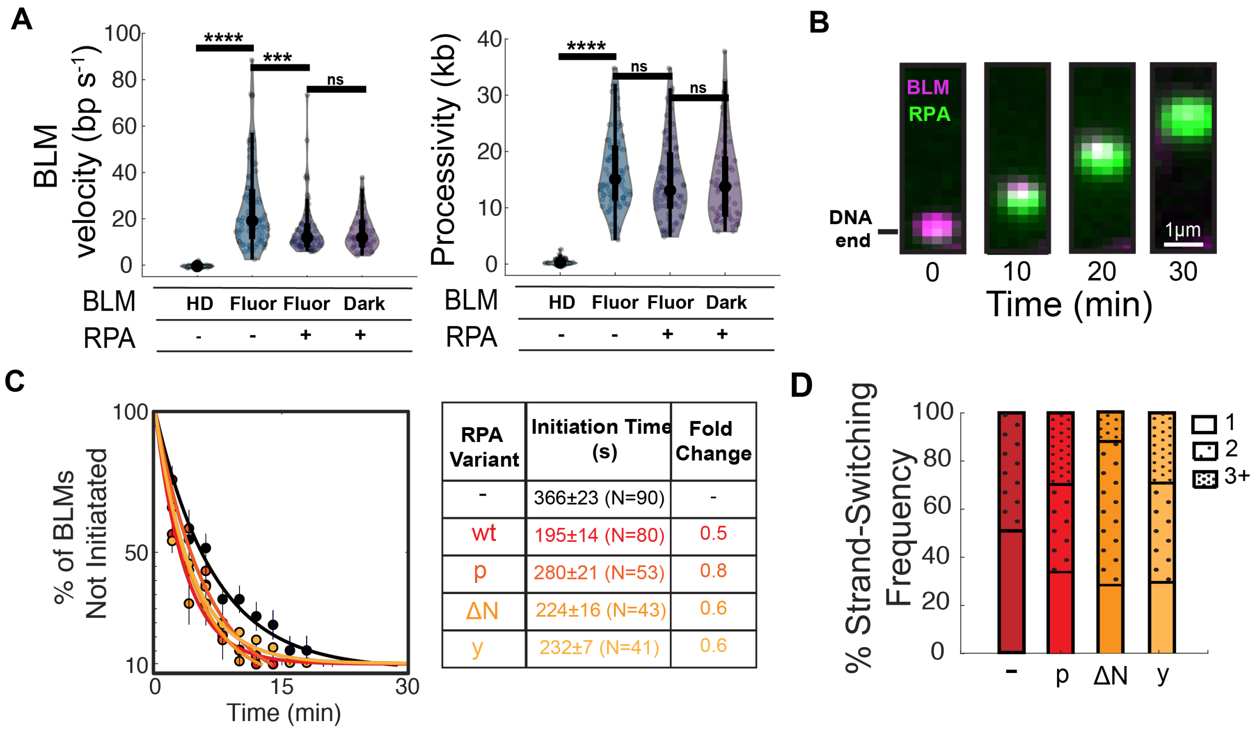


| RPA Variant | Initiation Time (s) | Fold Change |
|---|---|---|
| - | 366±23 (N=90) | - |
| wt | 195±14 (N=80) | 0.5 |
| p | 280±21 (N=53) | 0.8 |
| ΔN | 224±16 (N=43) | 0.6 |
| y | 232±7 (N=41) | 0.6 |

**Figure S3. Characterization of RPA regulation of BLM helicase**

**(A)** Velocity and processivity distributions showing fluorescent BLM and BLM without a fluorescent label are statistically indistinguishable. **(B)** RPA-GFP intensity increased with BLM helicase activity due to accumulation of ssDNA during BLM translocation. **(C)** BLM helicase initiation analysis with RPA variants. Error bars: S.D. as determined by bootstrap analysis. **(D)** pRPA increases the strand-switching frequency of BLM helicase.

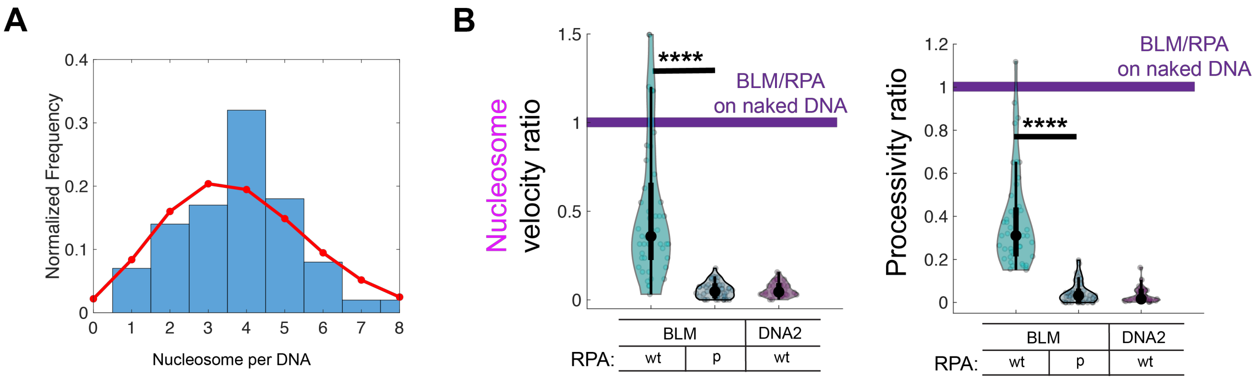


**Figure S4. Characterization of resection on nucleosome-coated DNA**

**(A)** Quantification of number of nucleosomes per DNA (N=100 DNA molecules analyzed). Red line represents a Poisson fit. **(B)** Relative velocity and processivity of fluorescently-labeled nucleosomes encountered by BLM/RPA or BLM/pRPA. DNA2 cannot encounter nucleosomes because it is stationary on 3'-ssDNA overhangs in the absence of BLM (see Fig. S1F and (Cejka et al., 2010; Nimonkar et al., 2011; Zhou et al., 2015)). The velocity and processivity for DNA2/RPA on naked DNA are therefore compared to BLM/DNA2/RPA on naked DNA showing the requirement of BLM for DNA2 resection.
